# Dynamics of a hepatocyte-cholangiocyte decision-making gene regulatory network during liver development and regeneration

**DOI:** 10.1101/2021.04.22.440352

**Authors:** Sarthak Sahoo, Ashutosh Mishra, Anna Mae Diehl, Mohit Kumar Jolly

## Abstract

Liver is one of the few organs with immense regenerative potential even at adulthood in mammals. It is composed of primarily two cell types: hepatocytes and cholangiocytes, that can trans-differentiate to one another either directly or through intermediate progenitor states, contributing to remarkable regenerative potential of the liver. However, the dynamical features of decision-making between these cell-fates during liver development and regeneration remains elusive. Here, we identify a core gene regulatory network comprising c/EBPα, TGFBR2 and SOX9 that underlies liver development and injury-induced reprogramming. Dynamic simulations for this network reveal its multistable nature, enabling three distinct cell states – hepatocytes, cholangiocytes and liver progenitor cells (hepatoblasts/oval cells) – and stochastic switching among them. Predicted expression signature for these three states are validated through multiple bulk and single-cell transcriptomic datasets collected across developmental stages and injury-induced liver repair. This network can also explain the experimentally observed spatial organisation of phenotypes in liver parenchyma and predict strategies for efficient cellular reprogramming among these cell-fates. Our analysis elucidates how the emergent multistable dynamics of underlying gene regulatory networks drive diverse cell-state decisions in liver development and regeneration.

## Introduction

The liver, the largest internal organ in the body, performs key physiological functions. It possesses remarkable regenerative ability and is capable of restoring its mass, architecture and function completely after injury (Gadd et al., 2020; Kopp et al., 2016). Both the major cell types seen in liver parenchyma – hepatocytes and cholangiocytes (biliary epithelial cells) – have been shown to be capable of dividing extensively (Kopp et al., 2016) and transdifferentiate into one another, thus contributing to liver regeneration (Deng et al., 2018; Schaub et al., 2018; Yanger et al., 2013).

In the liver, hepatocytes and cholangiocytes perform very different functions. While the former executes most metabolic functions, including bile secretion; the latter are biliary epithelial cells that line the bile duct tubules and serve to transport bile from liver to the small intestine. Developmentally, hepatocytes and cholangiocytes are both formed from a common progenitor cell type known as hepatoblasts. Hepatoblasts can co-express markers for both hepatocytes (HNF4α, CK18) and cholangiocytes (CK19) (Gordillo et al., 2015), a characteristic trait of such bipotent cells witnessed during embryonic development (Zhou and Huang, 2011). Cellular plasticity seen *in vivo* between cholangiocytes and hepatocytes suggests that while they may be terminally differentiated, they retain the ability to transdifferentiate into one another during injury-induced repair (Deng et al., 2018; Schaub et al., 2018). These observations suggest that these differentiated cells in liver can carry permissive chromatin from their progenitors, which may be important for their developmental and reprogramming competence (Li et al., 2020). However, the intracellular and tissue-level dynamics of cell-state transitions between these cell types in a liver remains unclear.

During chronic liver injury as well, cholangiocytes can transition to hepatocytes via an intermediate bi-phenotypic state which expresses both hepatocytic and biliary markers (Deng et al., 2018), reminiscent of earlier observations about the existence of oval cells – bipotent cells that expressed hepatoblast marker AFP and could differentiate to both lineages (Kopp et al., 2016). Similarly, SOX9^+^ hepatocytes have also been implicated to behave as bipotent progenitor cells after liver injury (Han et al., 2019; Tsuchiya and Yu, 2019). In liver homeostasis, liver progenitor cells (LPCs) have been identified that can differentiate into both hepatocytes and cholangiocytes in culture and can repopulate a liver after transplantation (Li et al., 2020). Thus, while molecular and functional similarities and differences in these different types of bipotent ‘hybrid’ cells identified are still being accrued, it is intriguing to note that across the contexts of embryonic development, homeostasis, liver injury and regeneration, one or more bipotent cell types have been identified.

Across these different contexts of liver development, injury repair and reprogramming, a variety of ‘master regulators’ have been identified that are capable of either inducing a hepatocyte cell-fate over cholangiocytes or *vice versa* or drive trans-differentiation of cholangiocytes to hepatocytes or *vice versa*. However, there are surprisingly few studies that attempt to uncover the fundamental traits of underlying gene regulatory networks formed by these ‘master regulators’ that govern cell fate transitions in liver development and regeneration.

Here, we first identify a core gene regulatory network involved in both developmental and regenerative cell-state transitions between hepatocytes and cholangiocytes and elucidate the dynamics of this network through a mechanism-based mathematical model. Our simulations suggest that this network is capable of enabling three states that can be mapped to hepatocytes, cholangiocytes and a progenitor-like phenotype. These model predictions are supported by analysis of diverse gene expression datasets collected during development and reprogramming scenarios. Further, these states were shown to be capable of transitioning to one another, under the influence of biological noise and/or external perturbations, deciphering possible mechanistic basis for observed trans-differentiation. We also decipher how the emergent properties of this regulatory network can give rise to spatial pattering of these liver cell types. Our mathematical model unravels design principles of the underlying liver cell-fate decision network as well as lays a predictive framework to investigate the dynamics of cell commitment and reprogramming in liver.

## Results

### Multistability in gene regulatory network underlying hepatocyte-cholangiocyte cell-fate commitment and plasticity

First, we identified a core gene regulatory network that is integral to the observed plasticity among hepatocytes, hepatoblasts and cholangiocytes. Various molecular players have been implicated in remarkable cellular plasticity of liver cells both in the context of liver development (Gérard et al., 2017; Lau et al., 2018; Zong and Stanger, 2011) and regeneration (Li et al., 2020). Among these, TGFβ signalling is of key importance in the differentiation of cholangiocytes from the hepatoblasts. High levels of TGFβ signalling is required near the portal vein for differentiation of biliary cells *in vivo* (Clotman et al., 2005). Consistently, TGFβ can mediate hepatocyte to cholangiocyte trans-differentiation *in vivo* (Schaub et al., 2018), and can suppress the transcription and activity of HNF4α, a well-known hepatocyte inducer (Cozzolino et al., 2013; Li et al., 2000; Lucas et al., 2004). Overexpression of TGFBR2 in hepatoblasts *in vitro* can drive them into a cholangiocyte fate (Takayama et al., 2014). Thus, TGFβ signaling can be thought of as a ‘master regulator’ of cholangiocyte cell-fate. Similarly, a ‘master regulator’ of hepatocyte cell fate is c/EBPα, whose overexpression transcriptionally inhibits TGFBR2 and can drive hepatocyte differentiation in hepatoblasts (Takayama et al., 2014). Furthermore, suppression of c/EBPα can stimulate biliary cell differentiation via increased Hnf6 and Hnf1b expression in periportal hepatoblasts (Yamasaki et al., 2006). In these hepatoblasts, overexpression of TGFBR2 inhibits c/EBPα, thus forming a ‘toggle switch’ or mutually inhibitory feedback loop between TGFBR2 and c/EBPα. Such ‘toggle switches’ are hallmarks of cellular decision-making between various ‘sibling’ cell fates (Zhou and Huang, 2011).

Another important player in biliary development is SOX9, which can repress the expression of both c/EBPα and TGFBR2 in mature biliary cells (Antoniou et al., 2009; O’Neill et al., 2014). SOX9, together with SOX4, can coordinate development of the bile duct (Poncy et al., 2015), during which it is upregulated by TGFβ signaling through the Jagged1-Notch axis (Wang et al., 2018). SOX9, similar to c/EBPα and TGFβ/TGFBR2 signaling, has been proposed to self-activate either directly or indirectly (Du et al., 2018; Duan and Derynck, 2019; Friedman, 2015). Put together, these interactions constitute a gene regulatory network comprising SOX9, c/EBPα and TGFBR2 (**Fig 1A**) which is involved in hepatocyte-cholangiocyte cell-fate decisions in the liver. Our goal here is not to identify a comprehensive network, but a minimal network motif that can potentially explain diverse instances of cell differentiation and reprogramming seen in liver development and injury repair. Thus, the nodes represented here – SOX9, c/EBPα and TGFBR2 – can be viewed as proxies for their corresponding co-factors and their regulons implicated in cell-fate commitment.

**Figure 1:**
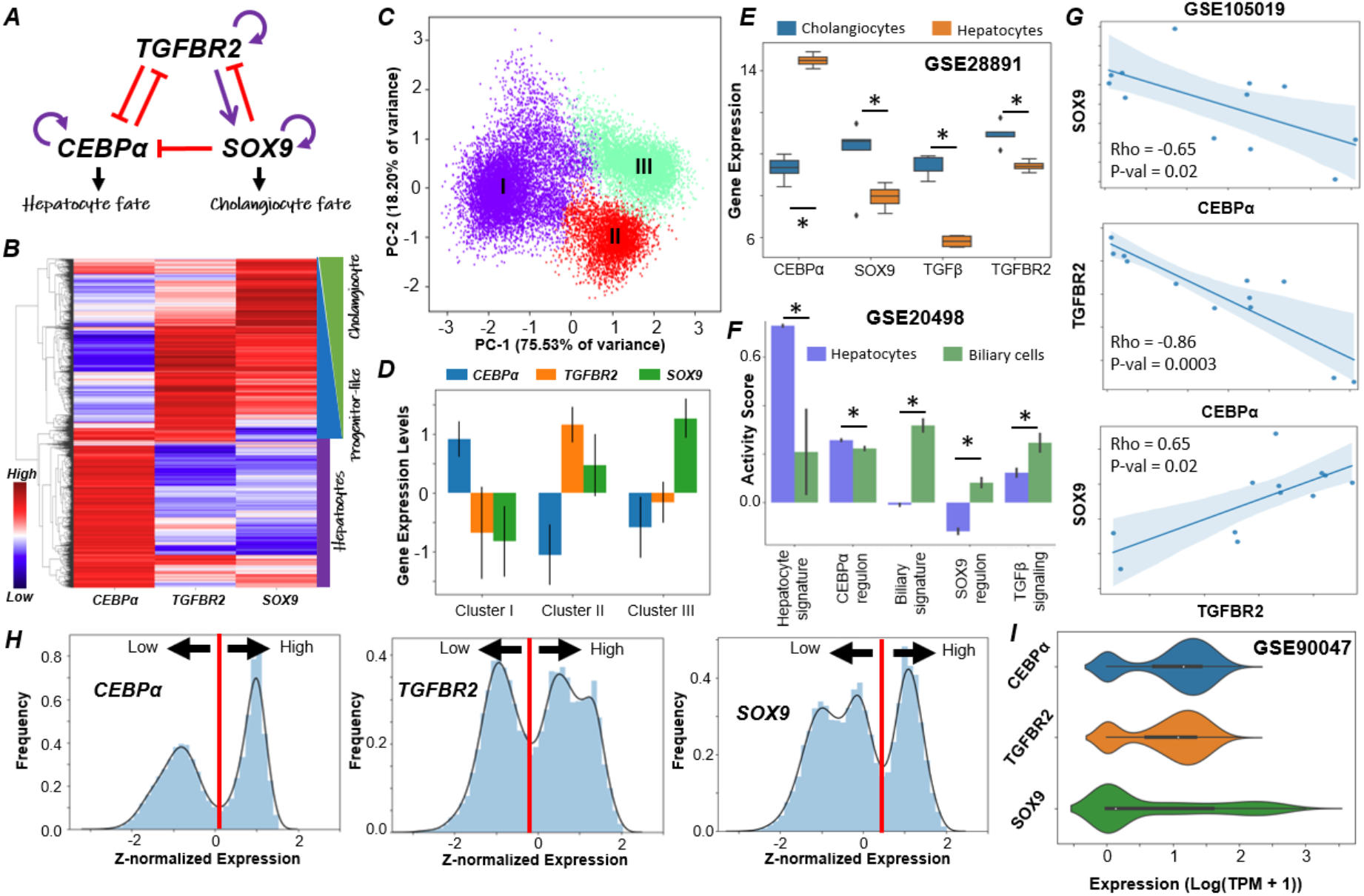
Multiple phenotypes enabled by a regulatory network driving cell-fate decisions in liver. **A)** Gene regulatory network (GRN) underlying plasticity in the hepatocyte-cholangiocyte cell-fate decision. Purple arrows represent activation links; red hammers represent inhibitory links. **B)** Heatmap showing hierarchical clustering in ensemble of steady state solutions obtained from the GRN. Red represents higher expression levels while blue denotes lower expression levels, plotted as z-scores. **C)** Principal Component Analysis (PCA) showing three major clusters in steady state solutions obtained. **D)** Quantification of steady state levels from the three clusters identified by PCA. Error bars represent the standard deviation over n=3 replicates. **E)** Gene expression levels of SOX9, CEBPα, TGFβ and TGFBR2 in hepatocytes and cholangiocytes (GSE28891). **F)** Activity levels of the gene expression signatures quantified through single sample Gene Set Enrichment Analysis (ssGSEA) in hepatocytes and biliary cells (GSE20498): CEBPα regulon, SOX9 regulon and TGFβ signalling. In E, F panels; * represents a statistically significant difference in the activity/expression levels (Students’ two tailed t-test; p-value <= 0.05) **G)** Scatter plots showing pairwise correlations in human liver progenitor-like population in culture (GSE105019). Spearman correlation coefficient (Rho) and p-value (P-val) are given. **H)** Simulation data indicating the multimodal nature of steady state values in the expression of the various nodes in the network. **I)** Single-cell RNA-seq data (GSE90047) for a population composed of hepatocytes, hepatoblasts and cholangiocytes: distributions of gene expression levels of SOX9, CEBPα and TGFBR2.

To understand the distinct phenotypes enabled by the gene regulatory network involving c/EBPα, TGFBR2 and SOX9 (**Fig 1A**), we performed dynamical simulations on this regulatory network, using an ensemble of kinetic parameter sets for a set of ordinary differential equations (ODEs) representing the regulatory interactions within a network. The parameter sets sampled randomly from a biologically plausible regime capture the inherent variability in a cell population. For each parameter set, various possible steady-state (phenotype) combinations are collated to identify the robust dynamical features emerging from this network topology (see **Methods**).

For the ensemble of steady-state solutions thus obtained, hierarchical clustering suggested the evidence of three distinct cell-states, through a heatmap (**Fig 1B**). A c/EBPα-high state can be attributed to a mature hepatocyte (Akai et al., 2014), whereas a SOX9-high state can be mapped on to a cholangiocyte profile (Dianat et al., 2014). Thus, progenitor-like (hepatoblasts or oval cells) state is likely to have low levels of both c/EBPα and SOX9, with possible enrichment of TGFBR2 (Takayama et al., 2014). Further, we observed a continuum between the progenitor-like and cholangiocyte phenotypes, pointing towards a gradient of cholangiocyte fate determination and maturation with possible intermediate states, as experimentally reported (Yang et al., 2017). Further credence for three states was obtained via principal component analysis (PCA), through which three distinct clusters of points were visually discernible (**Fig 1C**). The c/EBPα - high state (cluster I) had relatively low levels of SOX9 and TGFBR2. The TGFBR2-high state (cluster II) had moderate to low levels of SOX9 and low levels of c/EBPα. The SOX9-high state (cluster III) had low levels of both TGFBR2 and c/EBPα (**Fig 1D**). These observations were corroborated by experimental data from transcriptomic analysis of hepatocytes and cholangiocytes (GSE28891) (Shin et al., 2011) (**Fig 1E**). As predicted by our model, levels of c/EBPα was significantly higher in hepatocytes than in cholangiocytes, but those of SOX9, TGFβ and TGFBR2 were significantly higher in cholangiocytes than in hepatocytes (**Fig 1E**). In another dataset where the adult hepatocyte and cholangiocyte signatures were found to be significantly enriched in hepatocytes and cholangiocytes respectively, the c/EBPα regulon activity was higher in hepatocytes than in the biliary cell population while the SOX9 regulon and overall TGFβ signalling followed the opposite trend (GSE20498; **Fig 1F**) (Cullen et al., 2010). This analysis suggests that multistable dynamics of this gene regulatory network investigated here can allow for three distinct phenotypes – hepatocytes, cholangiocytes and bipotent progenitors (hepatoblasts/oval cells) – whose molecular footprints as predicted here are corroborated by diverse high-throughput transcriptomic datasets.

Next, we quantified pairwise correlation between these master regulators, and noted that while c/EBPα is negatively correlated with SOX9 and TGFBR2, SOX9 and TGFBR2 correlate positively (**Fig S1A**) in our simulations. This trend is consistent with observations in a population of primary hepatocytes and hepatocyte progenitor cells (**Fig 1G**; GSE105019) (Fu et al., 2019) and evidence during the hepatocyte-ductal trans-differentiation (O’Neill et al., 2014). Further, we plotted histograms for gene expression values of c/EBPα, SOX9 and TGFBR2 obtained from simulations, revealing their bimodality (**Fig 1H**). Such bimodality was validated from single cell RNA-seq performed specifically on a population consisting of hepatocytes, hepatoblasts and cholangiocytes (**Fig 1I**; GSE90047) (Yang et al., 2017).

The abovementioned observations are largely also seen in simulations of an expanded gene regulatory network that includes a separate node showing the TGFβ ligand itself (**Fig S1B-D**). Besides allowing for the three abovementioned phenotypes, this extended network also enabled another one with high levels of CEBPα and SOX9 (**Fig S1D**). This phenotype may correspond to bipotent SOX9^+^ hepatocytes as has been observed in various context of liver injury and regeneration (Han et al., 2019; Tanimizu et al., 2014). Put together, we demonstrate that the network motif involving c/EBPα, SOX9 and TGFBR2 can recapitulate various cell phenotypes observed in liver development and regeneration: hepatocytes, cholangiocytes and their bipotent progenitors.

### Temporal dynamics of cellular decision-making for hepatocytes and cholangiocytes during liver development

After investigating different possible phenotypes enabled by the network comprising c/EBPα, SOX9 and TGFBR2, we investigated the dynamics of cellular decision-making, particularly the role of TGFβ in controlling the transitions between hepatocyte and cholangiocyte cell states. For a set of kinetic parameters estimated from literature, we mapped different cell-states observed at various levels of TGFβ through a bifurcation (dose-response curve) diagram (see **Methods**). We found that at lower TGFβ levels, c/EBPα can exist at two levels – high and low – mapping onto a hepatocyte and a progenitor/cholangiocyte fate respectively. Correspondingly, when c/EBPα levels were high, TGFBR2 and SOX9 levels were both low. On the other hand, when TGFBR2 and SOX9 were high, the levels of CEBPα were lower, thus indicating bistability (**Fig 2A**). As levels of TGFβ increase, the levels of both TGFBR2 and SOX9 increase with a concurrent disappearance of the hepatocyte cell state (**Fig 2A**), which can be interpreted as the maturation of progenitor cells into a distinct cholangiocyte-like cell state and subsequent destabilisation of the hepatocyte cell fate at higher levels of TGFβ. In developmental contexts, it has been observed that the levels of TGFBR2 increase along with those of SOX9 as cholangiocytes develop/mature from an hepatoblast stage (Takayama et al., 2014), thus corroborating the bifurcation diagram seen here. This trend is also consistent with a graded mode in which cholangiocytes develop, i.e. not an abrupt change to a mature state (Yang et al., 2017).

**Figure 2:**
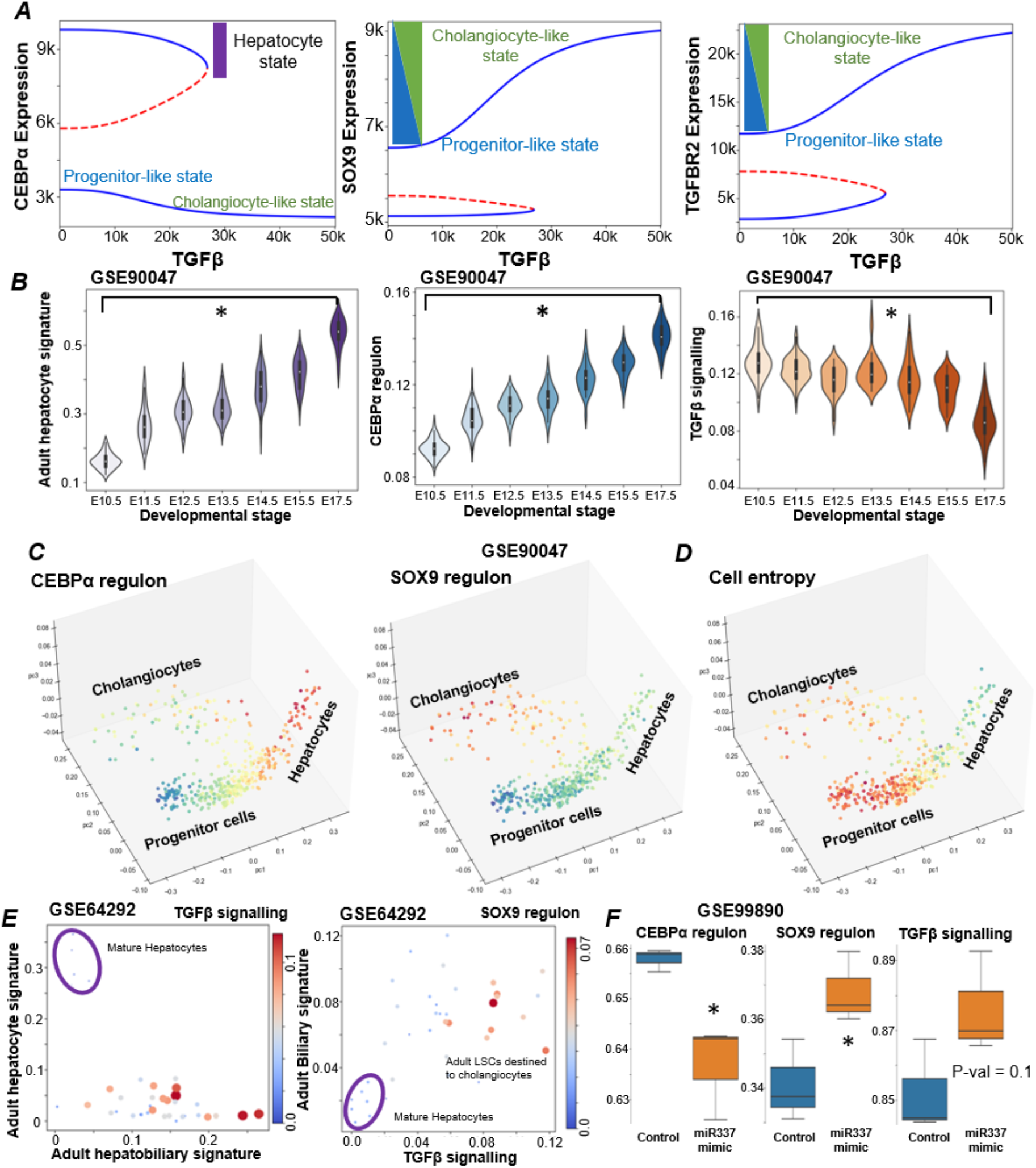
Dynamical signature of cell-state transitions in hepatocytes and cholangiocytes. **A)** Bifurcation analysis showing the stable (solid blue lines) and unstable steady state (dashed red lines) solutions in CEBPα, SOX9 and TGFBR2 levels, as a function of increasing TGFB levels. **B)** Experimentally observed adult hepatocyte signature, CEBPα regulon activity and TGFβ signalling activity in a population of single cells of hepatocytes/hepatoblasts at different developmental times (GSE90047). * represents a statistically significant difference in activity levels (Students’ two tailed t-test; p-value < 0.05) while comparing E10.5 and E17.5. **C)** Trajectory analysis of hepatocytes and cholangiocytes as they form from progenitor cells (coloured by CEBPα and SOX9 regulons showing their specificity to hepatocyte and cholangiocyte fate respectively). **D)** Cell entropy values seen to be the maximum in progenitor cells and decreases as cells enter more mature states. **E)** Single cell RNA-seq data (GSE 64292) on the hepatocyte – hepatobiliary signature plane coloured by TGFβ signalling activity (left) and SOX9 regulon activity (right). **F)** CEBPα regulon, SOX9 regulon and TGFβ signalling activity in hepatoblasts treated with miR-337 mimic showing a development towards cholangiocyte fate. * represents a statistically significant difference in the activity/expression levels (Students’ two tailed t-test; p-value <= 0.05)

To gain further confidence in these model predictions, we quantified the activity of CEBPα regulon and TGFβ signalling activity in a single-cell RNA-seq dataset (GSE90047) for sorted hepatoblasts, hepatocytes and cholangiocytes from E10.5-E17.5 mouse fetal livers state (Yang et al., 2017). As cells differentiated from hepatoblasts to hepatocytes, the adult hepatocyte signature and the c/EBPα regulon activity increased, while TGFβ signalling levels consistently decreased (**Fig 2B**). Similarly, as cells differentiated into cholangiocytes, activity for adult biliary signature and SOX9 regulon increase (**Fig S2A**). TGFβ signalling however showed an increased variance rather than a significant change in mean activity levels; one putative reason for which can be reduced levels of TGFBR2 in matured cholangiocytes (Antoniou et al., 2009). These observations were further corroborated by the same dataset, when we plotted the single-cell fate trajectory for data given in GSE90047 on a 3D PCA space, clearly showing the increase of c/EBPα levels in the hepatocyte branch and that of SOX9 higher in the cholangiocyte branch respectively (**Fig 2C**). However, TGFβ signalling, was active mostly in the cholangiocyte branch with intermediate levels of expression in the progenitor branch (**Fig S2B**) further underscoring the role of TGFβ signalling in commitment to the cholangiocyte cell fate. It was been earlier reported that progenitor cells generally have a larger transcriptional diversity (Gulati et al., 2020). To assess the transcriptional diversity in the cells, we computed the Shannon entropy values for individual cells (Teschendorff and Enver, 2017). Interestingly, we found out that entropy is the highest in the context of the progenitor cell and significantly lower in the hepatocytes (**Fig 2D**).

Similar expression patterns to those seen in liver development were observed in adult liver stem cells (GSE64292). Mature hepatocytes have very low levels of TGFβ signalling and SOX9 regulon activities (**Fig 2E**). Conversely, a subset of adult stem cells had high TGFβ signalling and SOX9 regulon activities, indicating a possible commitment towards the cholangiocyte fate (**Fig 2E**). Similarly, a sub-population of hepatoblasts expressing adult stem cell marker LGR5 have been shown to be capable of establishing both hepatocyte and cholangiocyte colonies (Prior et al., 2019). We found a bimodal distribution of SOX9 regulon and TGFβ signalling activity in the entire hepatoblast population as the developmental time progressed; however no such shift in the levels was observed in the context of LGR5+ hepatoblasts cells (**Fig S2C**), suggesting that a switch in TGFβ signalling and SOX9 levels are required to commit the hepatoblast cells towards a cholangiocyte or a hepatocyte cell fate. Previously, miR-337 has been implicated to promote a cholangiocyte fate during liver development (Demarez et al., 2018). We found that among differential gene expression programs induced by miR-337 mimic in *in vitro* cultures of immortalized hepatoblasts from E12.5 wild-type mice liver, c/EBPα regulon activity decreased significantly, but SOX9 regulon and TGFβ signalling increased (**Fig 2D**), indicating the functional role of these players in committing to a cholangiocyte cell-fate. Overall, this analysis highlights the role of interconnected emergent dynamics of c/EBPα, TGFBR2 and SOX9 in enabling different cell-fates during hepatic development and adult liver.

### Stochastic state switching and spatial pattern formation among liver cell phenotypes

After observing distinct cellular phenotypes enabled by this gene regulatory network, we examined whether these phenotypes could stochastically switch among one other under the influence of noise in gene expression. Previous experimental and computational studies have demonstrated the importance of intrinsic and extrinsic gene expression noise in cell fate switching in development and disease (Balázsi et al., 2011), including pathological conditions in liver (Sahoo et al., 2020a). We conducted stochastic simulations for the core gene regulatory network (**Fig 1A**) (Kohar and Lu, 2018) and observed switching among cell fates for various parametric combinations. We observed that hepatoblasts (characterised by high levels of TGFBR2 and low levels of both SOX9 and c/EBPα) could switch to a cholangiocyte cell fate (characterised by high level of SOX9) and stably exist for relatively longer periods of time (**Fig 3A** – top panel). Similarly, hepatocytes (high levels of c/EBPα) and cholangiocytes (high levels of SOX9) could switch between one another (**Fig 3A** – middle panel) indicative of trans-differentiation as observed in experimental studies (Deng et al., 2018; Schaub et al., 2018), We also observed parameter sets that showed dynamic transitions between hepatocytes, cholangiocytes, hepatoblasts and SOX9^+^ hepatocytes (characterised by intermediate levels of both SOX9 and c/EBPα) (**Fig 3A** – bottom panel). These results indicate that multistable features of this gene regulatory network can confer cellular plasticity in liver, which may be of paramount importance during injury repair and reprogramming.

**Figure 3:**
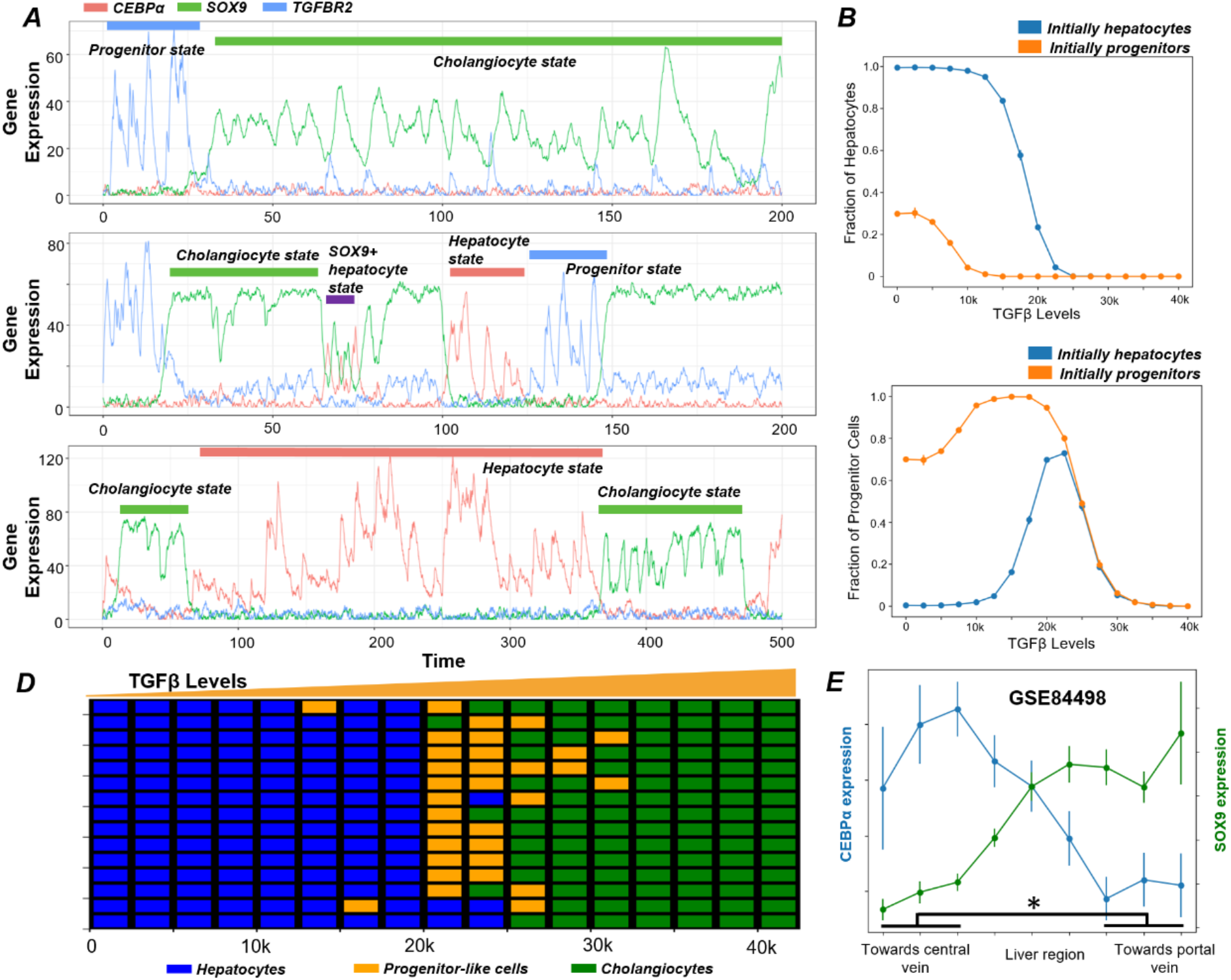
State switching and spatial patterns in liver. **A)** Noise induced stochastic switching simulations of the gene regulatory network (GRN) under different representative parameter sets. Biological states have been defined based on the expression levels of the individual genes. **B)** Effects of different initial conditions on the stochastic dynamics of GRN, showing hysteresis in the system – fraction of hepatocytes and fraction of progenitor cells. To test for hysteresis, two distinct initial conditions used for the system is either a pure hepatocyte or a progenitor-like population, and the fraction of the two cell populations are plotted. Error bars represent standard deviation values (n=3) **C)** Spatial patterning emergent from the dynamics of the gene regulatory network as a function of a gradient of TGFβ signalling under the influence of gene expression noise. **D)** Experimentally observed expression levels of c/EBPα and SOX9 via RNA-seq data (GS84498) in different regions of liver (towards central vein or portal vein). * represents a statistically significant difference in SOX9 and c/EBPα expression levels (Students’ two tailed t-test; p-value <= 0.05).

A hallmark feature of multistable regulatory networks is hysteresis. To characterize the hysteretic behaviour for this network, we first considered a “pure population” of hepatocytes and monitored, as a function of TGFβ levels, the proportion of cells that switched to the progenitor/cholangiocyte state and the proportion of cells that did not switch states. Similarly, we started from a population of “pure progenitors”, and tracked the relative proportion of cells in the three states (hepatocytes, cholangiocytes and progenitor cells) as a function of TGFβ levels. For a non-hysteretic system in nature, the proportions of cells in each of these three states would have been the same from the two starting populations (only hepatocytes or only progenitor cells) for a given level of TGFβ. However, we observed distinctly different profiles for the different starting populations (**Fig 3B**). For instance, at TGFβ=0, and starting from a population of only hepatocytes, the cells show minimal switching, instead maintain hepatocyte fate (**Fig 3B** – top panel). But if one starts from a population of only progenitor cells, at TGFβ=0, only ~30% switch to a hepatocyte fate and the remaining 70% retain their progenitor cell fate. Similarly, at TGFβ = 15,000 molecules, if we start from a population of only progenitor cells, almost none of them switch to a hepatocyte fate but if we start from a population of only hepatocytes ~20% of the cells will switch to a progenitor cell fate (**Fig 3B** – bottom panel). We also observed that the proportion of cholangiocytes obtained (when starting from a pure population of either hepatocytes or oval cells) has little effect on its starting population (**Fig S2E**), which is expected, given that the cholangiocyte state is stable mostly beyond the bistable region where the hysteretic pattern is less pronounced. While hysteresis in liver cell phenotypes remains to be validated, it has been experimentally witnessed for many multistable biological systems such as epithelial-mesenchymal plasticity and lactose metabolism (Celià-Terrassa et al., 2018; Ozbudak et al., 2004).

After examining the temporal dynamics of cell-fate commitment in the liver, we focused on spatial patterning of these phenotypes. Cholangiocytes are more abundantly found near the portal vein. On the contrary, hepatocytes are found much more abundantly on the parenchymal side (Clotman et al., 2005). Hepatic progenitor cells (similar to hepatoblasts/oval cells in our model) have been proposed to be sandwiched between the cholangiocytes and hepatocytes in the adult liver (Ko et al., 2020). Hepatoblasts in the periportal region are likely to adopt a cholangiocyte fate and eventually form the bile ducts, while hepatoblasts in other regions of the lobule form hepatocytes (Raynaud et al., 2011). Such spatial segregation has been proposed to be a result of TGFβ gradient (Ayabe et al., 2018; Clotman et al., 2005) and/or by mechanical cues induced Notch signalling variations in the periportal region (Kaylan et al., 2018).

We probed whether we could reproduce this spatial patterning of hepatocytes/hepatoblast-like cells/cholangiocytes by simulations capturing a simple gradient of TGFβ influence the gene regulatory network. Our simulations could produce the spatial pattern of phenotypes consistent with these observed trends. We demonstrated that under high levels of TGFβ, a cholangiocyte cell state are more prevalent, while at intermediate and low levels of TGFβ, hepatocytes and progenitor-like cells are seen to be abundant with the hepatocyte abundance increasing as the levels of TGFβ drop (**Fig 3D**). Our simulation observations are further corroborated by spatial gene expression analysis of liver tissue (GSE84498) (Halpern et al., 2017). We observed that the expression levels of c/EBPα drop towards the portal vein region with a concurrent increase in the levels of the SOX9 (**Fig 3E**). These observations provide insights into the spatial organisation of distinct cell types seen in liver development and homeostasis as a function of underlying gene regulatory networks.

### Cellular reprogramming strategies among liver cell phenotypes based on gene regulatory network dynamics

After elucidating the dynamics of a core gene regulatory network for cellular decision-making during liver development and homeostasis, we compared the reprogramming efficiencies of various perturbations on this regulatory network and the final proportions of cell fates obtained. We performed either over-expression (OE) or down-regulation (DE) individually on CEBPα, SOX9 and TGFBR2 and accessed the proportion of hepatocytes, cholangiocytes, progenitor-like cells and SOX9^+^ hepatocytes (see **Methods** for how the cell types have been defined for this analysis). Overall, we found that hepatocytes can be best enriched by either performing a CEBPα OE or TGFBR2 DE (**Fig 4A**). On the other hand, cholangiocytes can be best enriched for by SOX9 OE or TGFBR2 OE (**Fig 4B**). For the progenitor-like state, as it is characterised by high TGFBR2 levels and low levels of both CEBPα and SOX9, one might expect that over-expression of TGFBR2 levels would lead to an enrichment of this state. However, we find out that TGFBR2 OE has a weaker impact in enriching progenitor cell population as compared to CEBPα OE or SOX9 OE (**Fig 4C**). This observation lends support to the hypothesis that to maintain a progenitor cell state population, the differentiation programmes (both cholangiocyte-inducing and hepatocyte-inducing) need to be inactive. This observation is likely to be more generic, given that various underlying gene regulatory networks for cell-fate decisions have similar topology as the one shown here.

**Figure 4:**
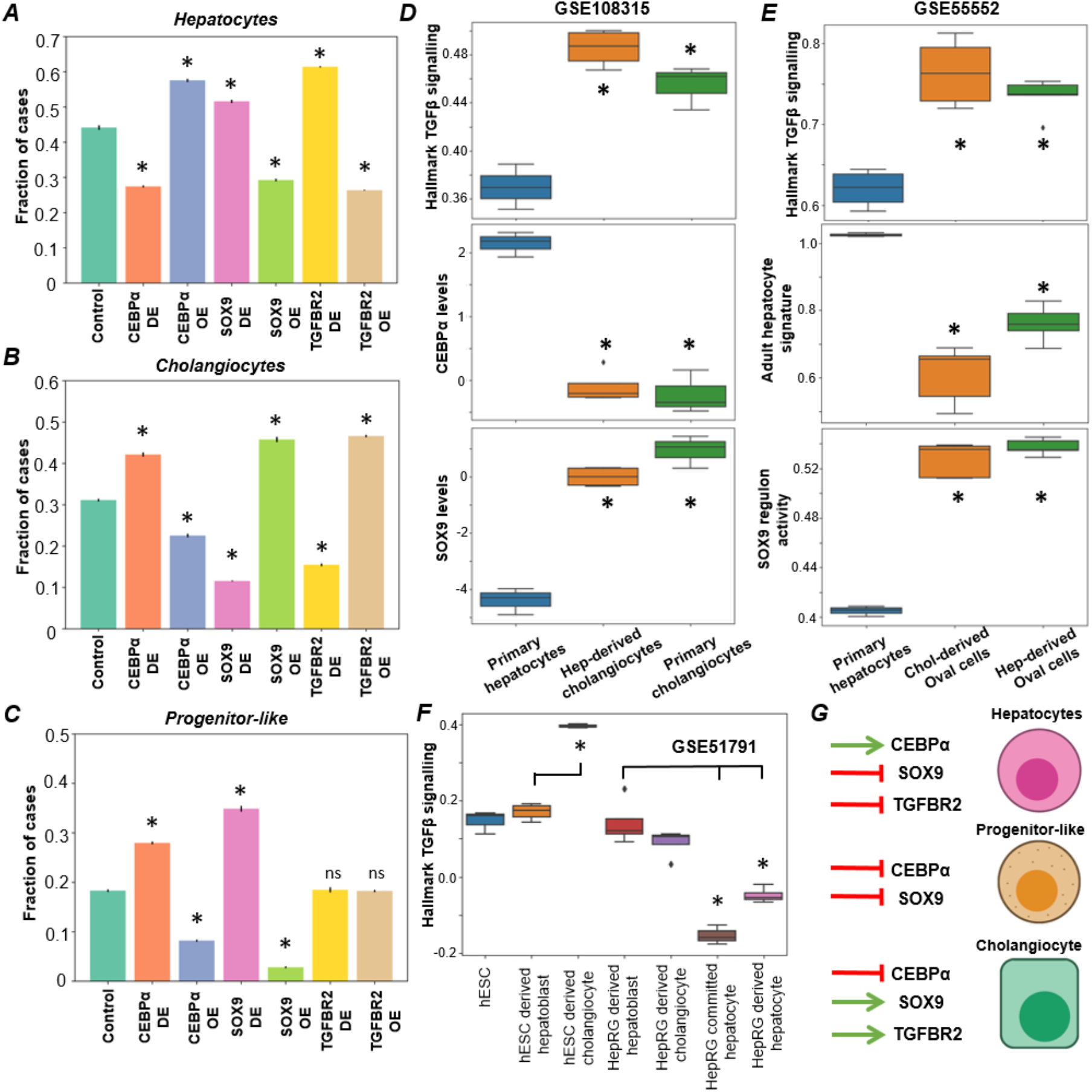
Predicted cellular reprogramming strategies. Simulation results quantifying the levels of **A)** hepatocytes **B)** cholangiocytes and **C)** progenitor-like cells under various perturbations to the gene regulatory network. * represents a statistically significant difference in the proportion of cases resulting in a particular phenotype compared to control case (Students’ two tailed t-test; p-value < 0.05); ns denotes a non-significant difference. **D)** Experimentally observed gene expression levels of CEBPα, SOX9 and activity of TGFβ signalling in a population of hepatocytes, cholangiocytes and cholangiocytes derived from hepatocytes (GSE108315). **E)** Experimentally observed activity levels of TGFβ signalling, hepatocyte signature and SOX9 regulon activity in hepatocytes and oval cells derived from either hepatocytes or cholangiocytes (GSE55552). **F)** Experimentally observed activity levels of TGFβ signalling in several reprogrammed cells (GSE51791). **G)** Schematic showing the list of perturbations that are likely to be the most effective in enriching for a given cell type. * represents a statistically significant difference in activity/expression levels with respect to the corresponding control cases (Students’ two tailed t-test; p-value < 0.05).

All the above-mentioned trends remain qualitatively unchanged when TGFβ was perturbed too. Further, SOX9^+^ hepatocytes, as expected, were most enriched for by SOX9 OE instead of by TGFBR2 OE, although SOX9 is activated by TGFβ signalling (**Fig S3A-B**), implying that enriching SOX9^+^ hepatocytes may need a more specific agonist facilitating SOX9 transcription, instead of a generic activation of TGFβ signalling pathway. Finally, expected correlation trends among CEBPα, SOX9 and adult signatures were witnessed in hepatic progenitor cell populations isolated from alcoholic steatohepatitis livers (GSE102683; **Fig S3C-D**) (Ceulemans et al., 2017).

Next, we assessed whether gene expression changes observed in hepatic development are also seen in the context of reprogrammed cells or in the context of trans-differentiated cells during liver injury. TGFβ treatment has been shown to convert hepatocytes to cholangiocytes via trans differentiation. In a peripheral bile duct RNA-seq dataset from mouse livers (GSE108315; **Fig 4D**), we observed TGFβ signalling to be higher in cholangiocytes and cholangiocytes derived from hepatocytes, as compared to low levels of TGFβ signalling in primary hepatocytes (Schaub et al., 2018). Similar trends are noted for SOX9, while CEBPα levels show the opposite trends, as expected (**Fig 4D**). In another instance, in mouse oval cells (progenitor-like cells during liver regeneration) either derived from cholangiocytes or hepatocytes through chronic injury (Tarlow et al., 2014), levels of SOX9 regulon and TGFβ signalling were higher than in primary hepatocytes, but the adult hepatocyte signature was lower (GSE55552; **Fig 4E**). Finally, during reprogramming from human embryonic stem cells (hESCs) and/or hepatic stem cell line (HepaRG) (Dianat et al., 2014), hepatocytes – both mature or immature – exhibit low levels of TGFβ signalling activity (compare 4^th^ column with the 6^th^ and 7^th^ columns in **Fig 4F**), while cholangiocyte had higher or comparable levels of TGFβ signalling activity when compared to hepatoblast cell state (compare 2^nd^ column with 3^rd^ column in **Fig 4F**), indicative of different levels of maturity of cholangiocytes (GSE51791). Together, this integrative analysis helps identify how modulating the relative levels of SOX9, CEBPα or TGFBR2 can drive diverse trajectories of cell state-switching across liver development, liver injury induced reprogramming and trans-differentiation scenarios (**Fig 4G**).

Besides OE or DE of nodes in the network, we also investigated possible trajectories of cellular reprogramming enabled by modulating the strength of individual edges in the network. Given that TGFBR2 DE was shown to enrich for hepatocytes, we probed the impact of a stronger inhibition of TGFBR2 by SOX9 (**Fig 5A**). For the control case, the bifurcation diagram showed that at TGFβ=0, the system is bistable (**Fig 2A;** blue curve in **Fig 5B**), i.e. in the absence of TGFβ signaling, a certain fraction of progenitor-like cells will be present in the population, thus blocking a transition to all-hepatocyte population, when starting from a population of only cholangiocytes. This need not necessarily be the case *in vivo* conditions. However, upon increasing the strength of suppression of TGFBR2 by SOX9, the bifurcation diagram changed, now enabling a scenario of only hepatocyte cells at TGFβ =0, with modest changes in steady state expression values of c/EBPα and SOX9 (green curve in **Fig 5B**). Further, cholangiocytes and hepatocytes can co-exist and likely stochastically switch even at higher values of TGFβ signaling, without necessarily requiring to transition via a hepatoblast stage. Thus, a stronger inhibition of TGFBR2 by SOX9 provides a permissive situation for the experimentally observed direct transdifferentiation among the two mature liver cell types (Deng et al., 2018; Schaub et al., 2018).

**Figure 5:**
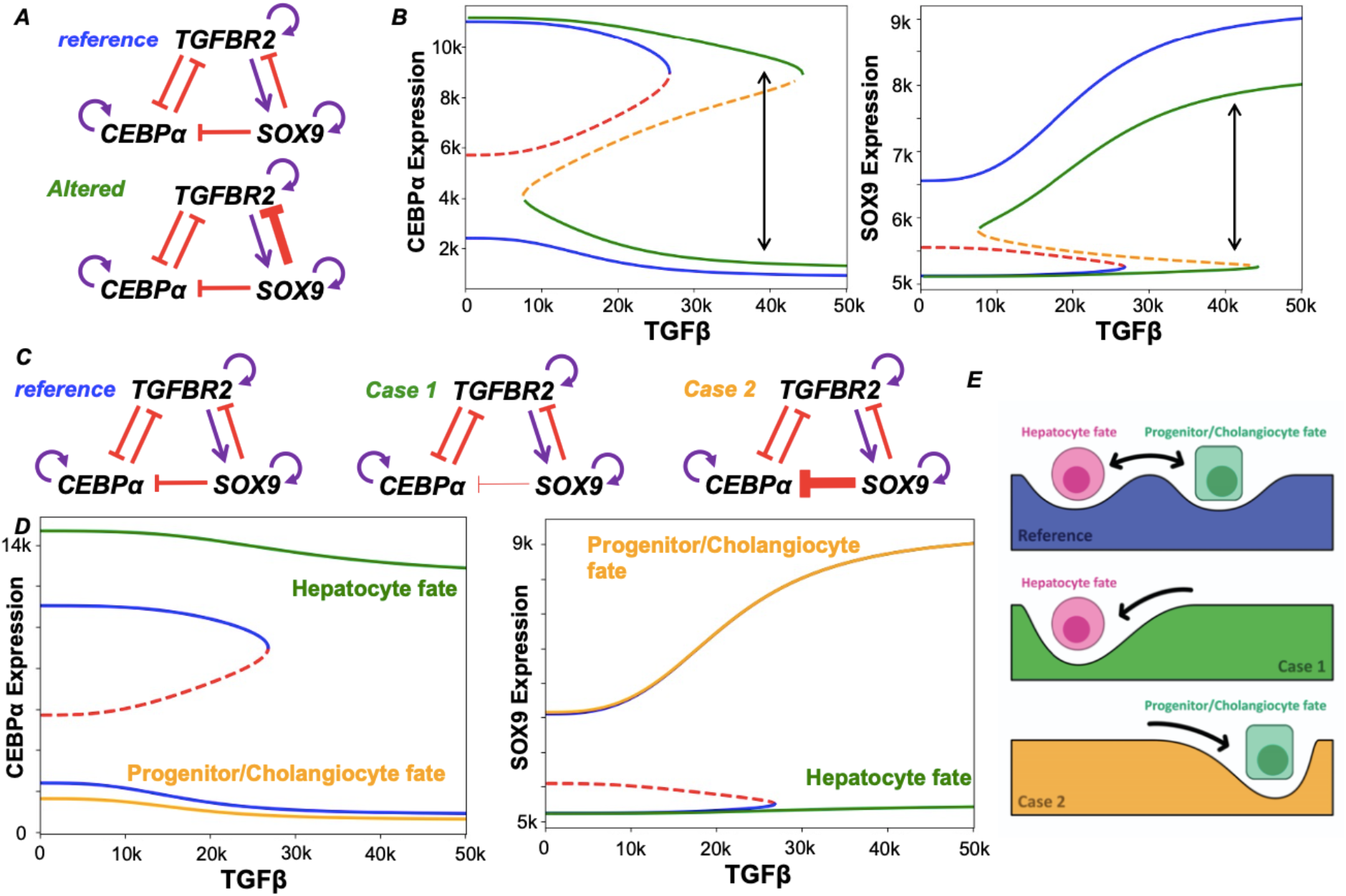
Impact of altering edge strengths on cell reprogramming landscape. **A)** Schematic showing the reference and the altered gene regulatory network in which the link strength of SOX9 inhibiting TGFBR2 has been increased. **B)** Changes in bifurcation diagram upon altering the circuit as shown in A. Blue solid lines show stable steady states while red dashed line shows the unstable states for the reference bifurcation diagram. Green solid lines and orange dashed lines are the stable and unstable states for the altered GRN in A. Bidirectional arrow is indicative of direct transition of hepatocytes to cholangiocytes at higher levels of TGFβ ligand. **C)** Schematic showing the reference and the two cases of the gene regulatory network in which the link strength of SOX9 inhibiting CEBPα has been decreased (Case 1) or increased (Case 2). **D)** Reference bifurcation diagram is shown in blue solid lines and red dashed lines. Bifurcation diagram for case 1 is shown in green solid line (hepatocyte state only) while the orange solid curve refers to case 2 (progenitor/ cholangiocyte state only). Note that there is no unstable states in the green and the orange profiles indicating that only one fate is enabled by the parameter set. **E)** Schematic showing the changes in the landscape brought about by changing the strength of SOX9 suppression of CEBPα in which for case 1 and case 2 only one stable state is allowed while for the reference case 2 stable states are possible.

Next, we probed the impact of altering the link strength from SOX9 to c/EBPα in both directions: a weaker inhibition (case I) or a stronger inhibition (case II) (**Fig 5C**). In case of weaker inhibition, the progenitor-like and cholangiocyte states are not observed irrespective of the value of TGFβ (green curve in **Fig 5D**). In contrast, in case of a stronger inhibition, hepatocyte state disappears (orange curve in **Fig 5D**). Thus, by changing the link strength from SOX9 to c/EBPα, landscape of cellular plasticity can be transformed such that only one cell state (‘attractor’) exists (**Fig 5E**), compromising on the ability to switch cell-fates in the liver. These observations suggest potent mechanisms to lock into specific cell-fates as well as pinpoint how a delicate balance between different links in underlying regulatory network allow multistability and consequent phenotypic plasticity – a fundamental tenet of development and regeneration across organs and organisms.

## DISCUSSION

Phenotypic plasticity is a ubiquitous phenomenon via which cells can exhibit diverse phenotypic states which can interconvert among themselves. It is thought to be a ‘bet-hedging’ strategy that can be implemented by cell populations to survive through various bottlenecks, such as drug-tolerant persisters seen ranging from microbial populations to cancer cells (Sahoo et al., 2020b; van Boxtel et al., 2017). Understanding the first principles behind cell-fate decision making in developmental systems can be paramount to elucidating mechanistic underpinnings of phenotypic plasticity in biological systems in general. Furthermore, such an understanding can propel rational strategies for reprogramming of cell types *in vitro* (Qian et al., 2018). The hepatocyte-cholangiocyte decision-making offers an ideal system, given their common progenitor (hepatoblasts) that can differentiate into these divergent cell-fate trajectories. Furthermore, once differentiated, these cell still retain the capacity to trans-differentiate to one another during liver injury (Gadd et al., 2020).

Here, we identified and analysed a minimalistic gene regulatory circuit that can, in principle, explain various cellular phenotypes observed in the liver and their cell-fate decision under the influence of specific signalling cues. This network comprising SOX9, TGFBR2 and c/EBPα is multistable in nature; enabling multiple co-existing phenotypes that can switch in presence of biological noise. The experimentally observed spatial patterning of these cell types in the liver can also be explained as an emergent property of this gene regulatory network. Finally, we explore possible perturbations in the gene regulatory network that can enrich for various cell phenotypes, driving reprogramming.

The *in silico* model constructed here, like all model systems, has limitations. First, the hepatocyte and cholangiocyte phenotypes are proxied by individual ‘master regulator’ transcription factors, which makes it tricky to identify various heterogeneous ‘micro-states’ such as Axin+ hepatocytes, Tert^High^ hepatocytes, hybrid periportal hepatocytes and Lgr5+ hepatoblasts (Li et al., 2020; Prior et al., 2019). Expanding the regulatory network to include other nodes may alleviate this limitation; for instance, upon including TGFβ, our simulations showed the emergence of SOX9^+^ hepatocytes.

These cells are reported to behave as bipotent progenitors after liver injury (Han et al., 2019; Tanimizu et al., 2017). Second, we only investigate the transcriptional control of these cell-fates; they can be affected by other layers of regulation such as epigenetics (Aloia, 2021). Future efforts to interrogate the impact of chromatin changes on cell-fate decisions (Jia et al., 2019) will be important. Third, our model cannot distinguish biological differences that might exist between developmental bipotent progenitors (hepatoblasts) and adult bipotent progenitors (oval cells). Within the scope of this study, these diverse biological entities are considered to be similar but they may be different in *in vivo* conditions. For instance, excessive activation of TGFβ signalling in the liver is associated with appearance of “hepatobiliary” features; i.e. “hybrid” cells co-expressing both hepatocyte and cholangiocyte markers (Raynaud et al., 2011). Another manifestation of “hybrid” cells may be those co-expressing SOX9 with HNF4α and HNF1α (Akai et al., 2014). A systematic approach to map the regulatory networks involved in cell decision making during liver development and injury repair can characterize various cell-fates observed experimentally.

Our analysis opens up a wide range of questions for which experiments can be designed to test out novel hypothesis. Based on the bifurcation analysis presented here and prior literature, TGFβ treatment is known to promote SOX9 levels by increasing activity of the Notch/TGFβ signalling (Wang et al., 2018), but it still remains to be seen if the prolonged treatment of TGFβ antagonists is able to push the mature/immature cholangiocytes into a progenitor-like/hepatocyte cell fate. Furthermore, even if the programme is reversible in the initial stages, is there a point of no return that prevents cholangiocytes from reverting back to hepatocytes/progenitor-like states, similar to those observed in other instances of phenotypic plasticity? (Tripathi et al., 2020) If yes, what mechanisms underlie a distinction between cell decision-making (reversible) and cell-fate ‘locking’ (irreversible). Juxtaposing stability vs. plasticity (switching ability) in a multistable system can offer new insights into the operating principles invoked during canalization as well as organ injury repair.

## Materials and Methods

### RACIPE simulations

Random Circuit Perturbation (RACIPE) is a computational framework that allows an extensive exploration of the dynamical properties of a gene regulatory network (Huang et al., 2017). Only the network topology is provided as an input to simulation framework, which is then modelled as a set of x ordinary differential equations (where x is the number of nodes in the gene regulatory network). The change in concentration of each node in the network depends on production rate of the node, the effect of regulatory links incident on the node (modelled as a shifted Hill’s function (Lu et al., 2013)) and the degradation rate of the node. Each parameter in the set of unknown parameters for ordinary differential equations (ODEs) is randomly sampled from a biologically relevant range. After such sampling, the set of parameterised ODEs is solved to get different possible steady state solutions. The set of ODEs can be multistable, i.e. multiple sets of steady state concentrations satisfy the set of ODEs. The program samples 10000 different sets of parameters. For each parameter set, RACIPE chooses a random set of initial conditions (n = 100) for each node in the network and solves, using Euler’s method, with the set of coupled ODEs that represent interactions among the different nodes in a network. For each given parameter set and initial conditions, RACIPE reports the steady-state values for each of the nodes in the network. The steady state values were then Z-normalised where the z-normalized expression value (z_i_) is given by the term:

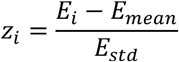

Where E_mean_ and E_std_ is the mean and standard deviation of the expression levels of a given node across all its steady state solutions.

The above procedure was followed to generate RACIPE results for the network topology given in **Fig 1A**. The Z-normalised steady state values were then plotted as a clustermap as in **Fig 1C**. PCA was performed with the major clusters being labelled via hierarchical clustering setting n = 3 as in **Fig 1B**. Kernel density maps overlaid on top of histograms for steady state expression for each node was plotted as in **Fig 1H**. Scatter plots for steady state solutions for each pair of nodes was plotted as presented in **Fig S1A**. The variant circuit as shown in **Fig S1B** was also simulated and processed using the methods described above to generate **Fig S1C** and **Fig S1D**.

The perturbation analysis in **Fig 4A-C** and **Fig S3A** were done by performing RACIPE analysis on the system by either over expressing (OE) or down expressing (DE) the given node by 10-fold. The Z-score normalisation of these perturbation data was done with respect to the control case. Similarly perturbation analysis was done on variant circuit shown in **Fig S1B** (shown in **Fig S3B**).

### Bifurcation analysis

We simulated a system of coupled ordinary differential equations (ODEs) using PyDSTool (Clewley, 2012) to create the bifurcations diagrams in the manuscript. The following set of ODEs were simulated:

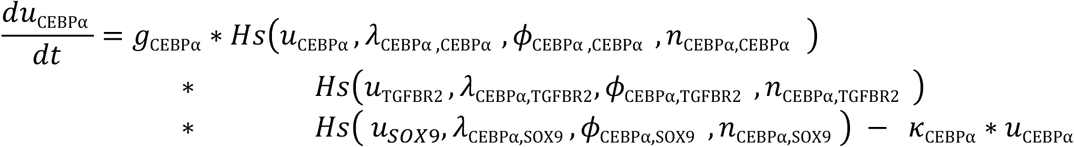

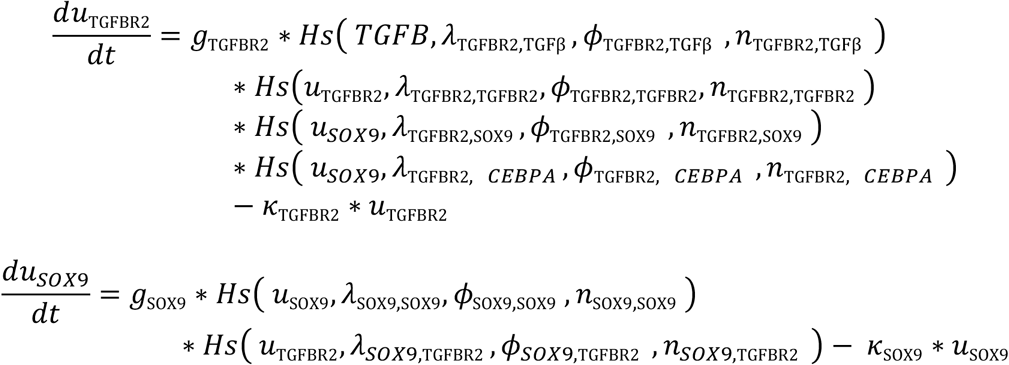

where g_i_ is the the production rate of the gene i, *k*_*i*_ is the degradation rate of the gene i, Hs is the shifted hills function that is given by the term: 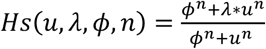

Each edge in the network has an associated shifted hills function which captures the effect of pair of nodes, i and j (j affecting i). *λ*_*i,j*_ is the fold change from the basal synthesis rate of i due to j. Therefore, λ > 1 for activatory links and λ < 1 for inhibitory links. *ϕ*_*i,j*_ is the threshold value for the interaction and *n*_*i,j*_ is the hills coefficient. The initial conditions for CEBPα, SOX9 and TGFBR2 were set to be 12000, 120 and 120 respectively. TGFβ levels was varied from 0 to 50000 units to create the bifurcation diagram. The parameters for the reference bifurcation diagram have been listed in **Supplementary Table 1**. Simulation of this system resulted in **Fig 2A**. For creating the bifurcations in **Figure 5B**, the parameter *λ*_TGFBR2,SOX9_ was changed from 0.5 (reference circuit) to (altered circuit). Similarly for the bifurcations in **Figure 5D**, the parameter *λ*_CEBPα,SOX9_ was varied from 0.5 to 0.65 (Case 1) and to 0.35 (Case 2).

### Stochastic simulations (including noise) – Hysteresis and Spatial dynamics

For the stochastic simulations in **Fig 3A** we used the webserver facility of Gene Circuit Explorer (GeneEx) to simulate stochastic dynamics of gene regulatory circuit as shown in **Fig 1A** — https://shinyapps.jax.org/5c965c4b284ca029b4aa98483f3da3c5/

For the simulating the hysteresis in the system we simulated the above described set of ODEs and with the parameter sets listed in **Supplementary Table 1** in the presence of noise at different values of TGFβ (0 to 40000 in steps of 2500) which was incorporated as follows:

1. We used Wiener process to simulate the noise because it a continuous gaussian white noise.
2. Wiener process is a continuous noise with the properties that

1. W(t = 0) = 0
2. The increments in W are gaussian and independent
3. For our model we have used a diagonal noise

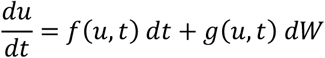

Here *f*(*u, t*) represents the deterministic equation and *g*(*u, t*) is the coefficient of the noise, and *dW* represents the Wiener process. For our case *g*(*u, t*) is a diagonal matrix, so each of the differential equations for CEBPα, TGFBR2, and SOX9 have their individual noise terms. For the simulation we used the noise values (*w*_1_, *w*_2_, *w*_3_) of 1000, 1000, 10 respectively.

The system was started from either a hepatocyte state (11000, 2500, 5100) or a progenitor-like state (2100, 12500, 6500) where the values are in the order of CEBPα, TGFBR2, and SOX9 respectively. A cell was deemed to be hepatocyte if the CEBPα level was above 6200 while it was deemed a progenitor like cell if the level of TGFBR2 was between 10000 and 17500 and values above that were labelled to be cholangiocytes. This analysis was carried out for a 3 sets of replicates each with 500 instances to estimate the mean and standard deviation for the fraction of cases for a given cell type.

For spatial simulations, we started the system from an all hepatocyte state and simulated 15 cells at each level of TGFβ (0 to 40000 in steps of 2500). Same classification of biological phenotypes as above was used to assign cell types on the spatial pattern.

### Data Analysis

All bulk microarray and RNA-Seq pre-processed datasets were obtained from publicly available GEO datasets. Gene signatures for adult hepatocyte programme, adult biliary programme and adult hepatobiliary programme were obtained from (Segal et al., 2019). The gene sets specifying CEBPα regulon and SOX9 regulon were obtained from (Møller and Natarajan, 2020). Hallmark TGFβ signalling pathway was obtained from MSigDB (Liberzon et al., 2011). Gene set activity were estimated via ssGSEA for all bulk samples and via AUCell for all single cell datasets (Aibar et al., 2017; Subramanian et al., 2005). For trajectory analysis the coordinates of each cell was obtained by performing a 3D PCA on the gene sets provided in (Segal et al., 2019).

### Statistical testing

We computed the Spearman correlation coefficients and used corresponding p-values to gauge the strength of correlations. For statistical comparison between groups, we used a two-tailed Student’s t-test under the assumption of unequal variances and computed significance.

## Author contributions

MKJ conceived and supervised research; SS and AM performed research; SS and AMD analysed data; SS prepared the first draft of the manuscript; all authors edited and wrote the manuscript.

## Acknowledgements

MKJ was supported by Ramanujan Fellowship awarded by Science and Engineering Research Board (SERB), Department of Science and Technology, Government of India (SB/S2/RJN-049/2018), and by InfoSys Foundation, Bangalore.

## Conflict of interest

The authors declare no conflict of interest.

## Data and code availability

All codes used in the manuscript are available at https://github.com/sarthak-sahoo-0710/2021_liver_cell_plasticity

All data used was through publicly available gene expression datasets, the GEO IDs for which are provided.

## Supplementary Figures

**Supplementary Figure 1:**
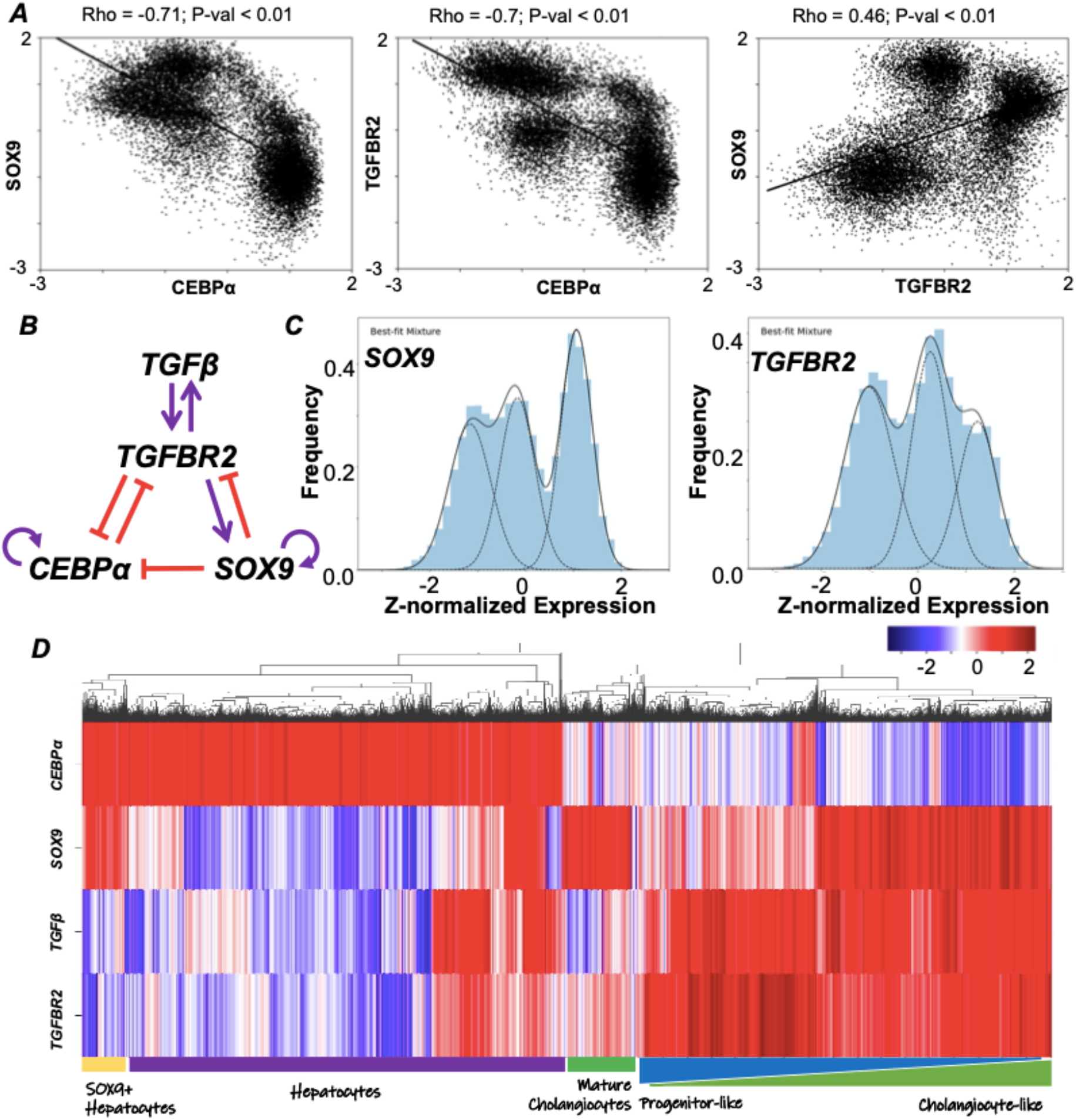
**A)** Scatter plots showing pairwise correlation between steady state values obtained via simulations for the given gene regulatory network in Fig 1A. Spearman correlation coefficient (Rho) and p-value (P-val) are given. **B)** A variant gene regulatory network showing TGFβ ligand explicitly (instead of it being lumped together with TGFBR2 as a single node). **C)** Multimodal gene expression levels of SOX9 and TGFBR2 for the simulated variant circuit. **D)** Cluster map showing the steady state solutions of the four nodes. The possible biological mapping of 3 distinct phenotypes have been labelled – hepatocytes, SOX9^+^ hepatocytes, and mature cholangiocytes. A possible continuum of progenitor-like cells to cholangiocyte-like cells is also shown. Red represents higher expression levels while blue denotes lower expression levels.

**Supplementary Figure S2:**
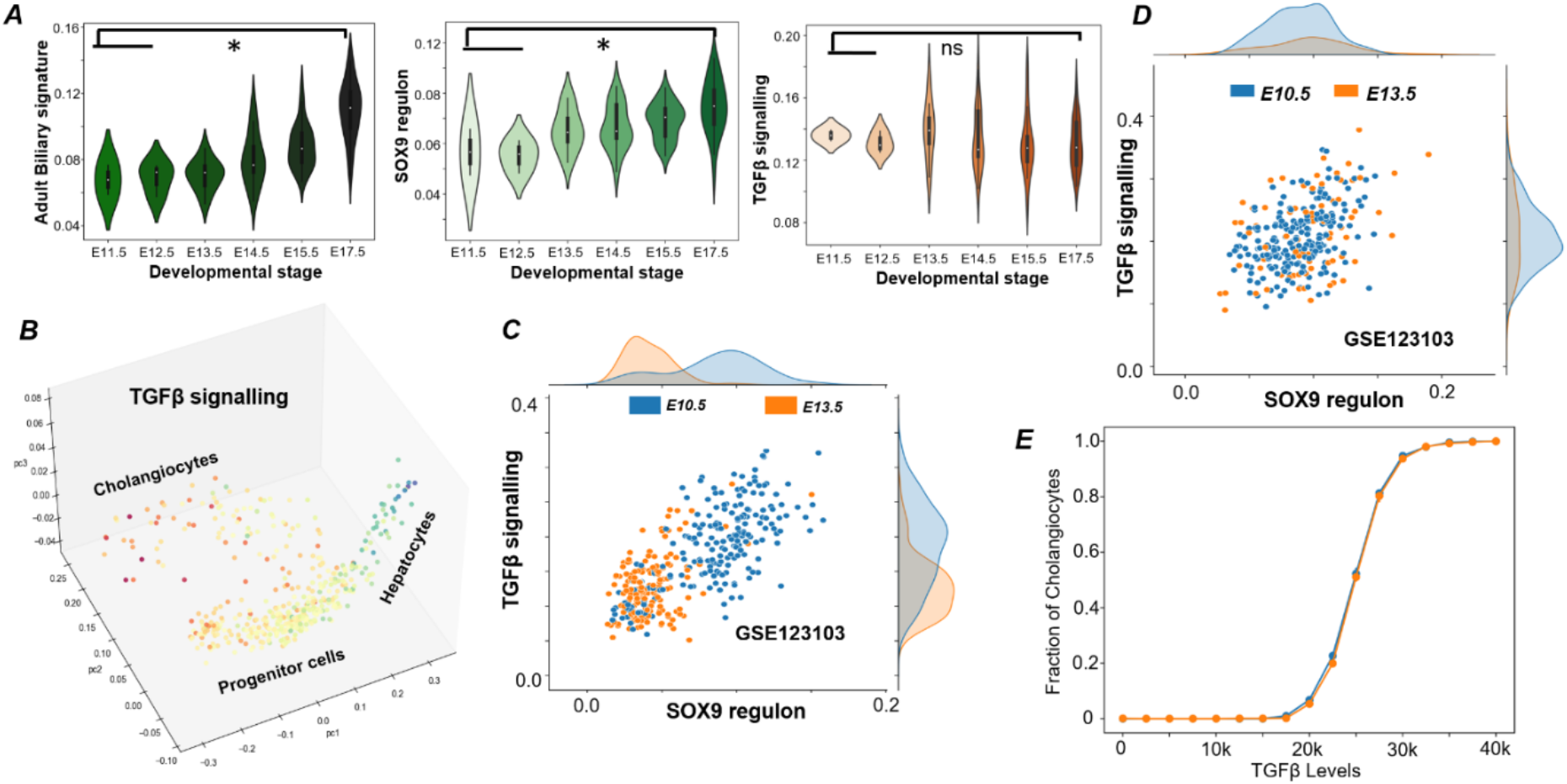
**A)** Experimentally observed adult biliary signature, SOX9 regulon activity and TGFβ signalling activity in a single-cell RNA-seq dataset containing mature/immature cholangiocytes with developmental time (GSE90047). * represents a statistically significant difference in activity/expression levels (Students’ two tailed t-test; p-value <= 0.05); ns indicated non-significant. **B)** Trajectory analysis of hepatocytes and cholangiocytes as they are formed from progenitor cells (coloured by TGFβ signalling activity). Note the relatively high/medium levels of TGFβ signalling activity in the cholangiocyte and the progenitor cell branch and its low expression in the hepatocyte branch. **C)** Scatter plot showing the TGFβ signalling activity and SOX9 regulon activity in a population of all hepatoblasts (LGR5+ or otherwise) at two distinct developmental stages. Note the relative drop in the values of both the quantities at E13.5. **D)** Scatter plot showing the TGFβ signalling activity and SOX9 regulon activity in a population of LGR5+ hepatoblasts only at the two developmental stages. **E)** Effects of different initial conditions on the stochastic dynamics of GRN, showing hysteresis in the system – fraction of cholangiocytes. To test for hysteresis, two distinct initial conditions used for the system is either a pure hepatocyte or a progenitor-like population, and the steady-state fraction of two cell populations are plotted. Error bars represent standard deviation values (n=3).

**Supplementary Figure S3:**
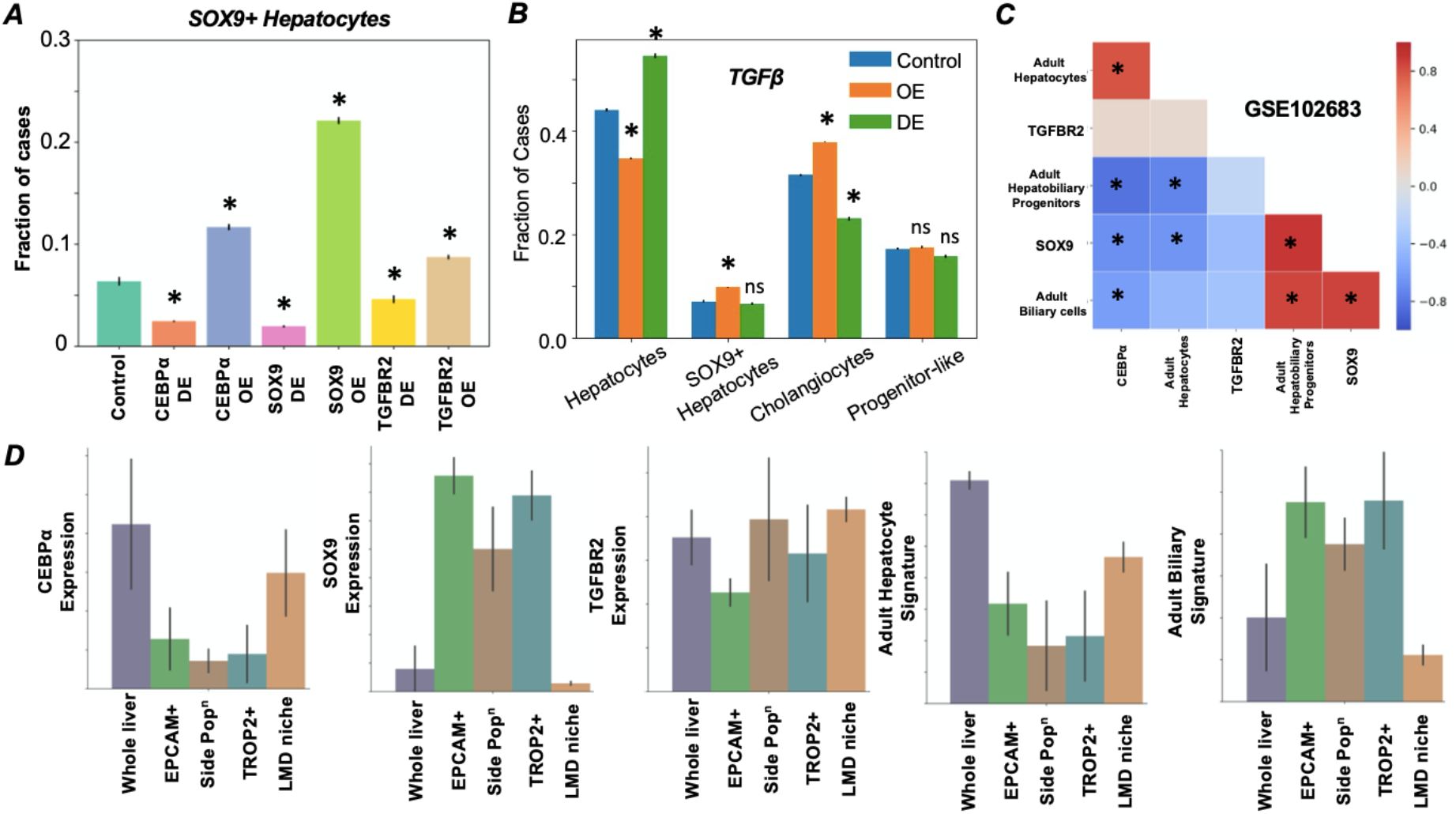
**A)** Simulation results quantifying the levels of SOX9^+^ hepatocytes under various perturbations to the gene regulatory network. * represents a significant difference in the proportion of cases resulting in a particular phenotype compared to the control case (p-value < 0.05) **B)** Simulation results quantifying the effects of perturbations to TGFβ in the variant circuit and the corresponding effects on the four phenotypes. * represents a significant difference in the proportion of cases resulting in a particular phenotype compared to the control case (p-value < 0.05); ns signifies a non-significant difference. **C)** Diagonal correlation matrix between expression levels (CEBPα, SOX9 and TGFBR2) and the gene expression signatures (adult hepatocyte, adult hepatobiliary progenitors and adult cholangiocytes) in bulk liver tissue samples (GSE102683). **D)** Quantification of CEBPα, SOX9, TGFBR2 expression levels along with activity of adult hepatocyte and adult biliary signatures in whole liver, EPCAM^+^, TROP2^+^, side cell populations and laser micro-dissected niche of adult liver stem cells via bulk RNA sequencing (GSE102683).

## Notes

### Competing Interest Statement

The authors have declared no competing interest.

